# Low-frequency variation in *TP53* has large effects on head circumference and intracranial volume

**DOI:** 10.1101/349845

**Authors:** Simon Haworth, Chin Yang Shapland, Caroline Hayward, Bram P. Prins, Janine F. Felix, Carolina Medina-Gomez, Fernando Rivadeneira, Carol Wang, Tarunveer S Ahluwalia, Martine Vrijheid, Mònic Guxens, Jordi Sunyer, Ioanna Tachmazidou, Klaudia Walter, Valentina Iotchkova, Andrew Jackson, Louise Cleal, Jennifer Huffmann, Josine L. Min, Lærke Sass, Paul R. H. J. Timmers, UK10K consortium, George Davey Smith, Simon E. Fisher, James F. Wilson, Tim J. Cole, Dietmar Fernandez-Orth, Klaus Bønnelykke, Hans Bisgaard, Craig E. Pennell, Vincent W. V. Jaddoe, George Dedoussis, Nicholas Timpson, Eleftheria Zeggini, Veronique Vitart, Beate Pourcain

## Abstract

Cranial growth and development affects the closely related traits of head circumference (HC) and intracranial volume (ICV). Here we model the developmental genetic architecture of HC, showing this is genetically stable and correlated with genetic determinants of ICV. Investigating up to 46,000 children and adults of European descent, we identify association with final HC and/or final ICV+HC at 9 novel common and low-frequency loci, illustrating that genetic variation from a wide allele frequency spectrum contributes to cranial growth. The largest effects are reported for low-frequency variants within *TP53*, with 0.5 cm wider heads in increaser-allele carriers versus non-carriers during mid-childhood.

## Main

The size and shape of the vertebrate brain is governed by the internal dimensions of the skull. Across vertebrate evolutionary history, major changes to brain size and proportion have been accompanied by modifications to skull morphology [1, 2]. This is also true within the lifespan of an individual, where developmental changes in brain size and shape must be reflected in changing cranial phenotypes.

Serial measures of maximal head circumference (HC) or occipito-frontal circumference (OFC) are routinely obtained to monitor children’s cranial growth and brain development during the first years of life and abnormal trajectories may indicate a range of neurological conditions [3]. In infants and children, HC is highly correlated with brain volume as measured by MRI studies [4, 5], especially in 1.7 to 6 year old children, although its predictive accuracy decreases with progressing age [5]. Healthy children from around the world, who are raised in healthy environments and follow recommended feeding practices, have strikingly similar patterns of growth [6]. The observation that final HC is largely determined by the age of 6 years in a large study from the UK [7] is therefore likely to be valid in multiple populations. In addition, nutritional status, body size and HC are closely correlated for healthy children during early life, and become less related after 24 months of age [8]. While HC properties in early childhood have immediate medical relevance, there are also compelling reasons to study HC in adulthood. In the adult population skeletal measures continue to act as a permanent measure of peak brain size that is unaffected by subsequent atrophic brain changes [9]. In early childhood, HC is likely to proxy overall body size and timing of growth, tracking changes in brain size. In older individuals, HC is valuable precisely because HC is robust to soft tissue atrophy, solely reflecting an absolute measure of final HC dimension.

HC is highly heritable and the notion of a developmentally changing, but etiologically interrelated, phenotypic expression of HC during the life course is supported by twin studies [10]. Reported twin-h^2^ estimates of 90% in infants, 85–88% in early childhood, 83–87% in adolescence and 75% in young and mid adulthood [10], with evidence for strong genetic stability between mid-childhood and early adulthood [10]. There are arguments to support the hypothesis that some of the underlying genetic factors act by a coordinated integration of signaling pathways regulating both brain and skull morphogenesis during development [11]. Especially, cells of early brain and skull are sensitive to similar signaling families [11]. Genetic underpinning of potentially shared mechanisms is supported by the fact that genome-wide signals for both infant HC and ICV are strengthened when combined [12], irrespective of their dissimilar developmental stages. However, genetic investigations studying (near) final HC and adult ICV are likely to be more informative on mechanisms underlying developmentally shared growth patterning which affect final cranial dimension. Additionally, low-frequency genetic variants, ranging between 0.5% to 5% minor allele frequency, have been poorly characterized by previous genome-wide association study (GWAS) efforts [13], both due to the small size of previous studies, and the limited coverage of lower-frequency markers by the first imputation panels.

Exploiting whole-genome sequence data together with high-density imputation panels such as the joint UK10K and 1000 genomes (UK10K/1KGP) [14] and the haplotype reference consortium (HRC) [15], that have previously facilitated the discovery of low-frequency genetic variants for a range of traits [16, 17], we carried out GWAS for final HC. Specifically, we aim to

a. study low-frequency and common variants for final HC, allowing for age-specific effects through meta-analyses of mid-childhood and/or adulthood datasets,
b. investigate genetic variants influencing a combined phenotype of (near) final HC and ICV, termed final cranial dimension, and
c. explore developmental changes in the genetic architecture of HC through longitudinal modelling of genetic variances in unrelated individuals as well as growth curve modelling of HC trajectories for carriers and non-carriers of high risk variants.

## Results

### Genome-wide analysis of HC scores

We carried out genome-wide analysis of HC scores using a two-stage developmentally-sensitive design (Figure 1a) including (i) pediatric (6 to 9 years of age), (ii) adult (16 to 98 years) and (iii) combined pediatric and adult samples comprising up to 18,881 individuals of European origin from 11 population-based cohorts and 10 million imputed or sequenced genotypes (Supplementary Table S1). In inverse-variance weighted meta-analysis (Supplementary Table S2-S4, Figure 2a-c, Supplementary Figure S1-S3), we identified three novel regions at chromosome 4q28.1 (HC (Pediatric): lead variant rs183336048, effect allele frequency (EAF)=0.02, *p*=3.0×10^−8^, Supplementary Figure S5a, S6a), 6p21.32 (HC (Pediatric)/ HC (Pediatric+adult): lead variant rs9268812, EAF=0.35, *p*=2.2×10^−9^, Supplementary Figure S5b, S6b) and 17p 13. 1 (Pediatric+adult: lead variant rs35850753, EAF=0.02, *p*=2.0×10^−8^, Supplementary Figure S5c, S6c, Figure 3a and b) as associated with HC at an adjusted genome-wide significant level (*p*<3.3×10^−8^) (Table 1). We followed-up the two signals in HC (Pediatric+adult) in a further 973 adults of European descent (mean age 50 years) (Supplementary Table S5, Supplementary Figure S6b,c) and replicated directionally consistent evidence for association with rs35850753 at the 17p13.1 locus (*p*=4.5×10^−5^, Table 1). In the combined pediatric, adult and follow-up sample, we observed here an increase of 0.24 sex-adjusted SD units in HC per increase in minor T risk allele (*p*=2.1×10^−10^, Table 1, Figure 3a and b).

**Figure 1.**
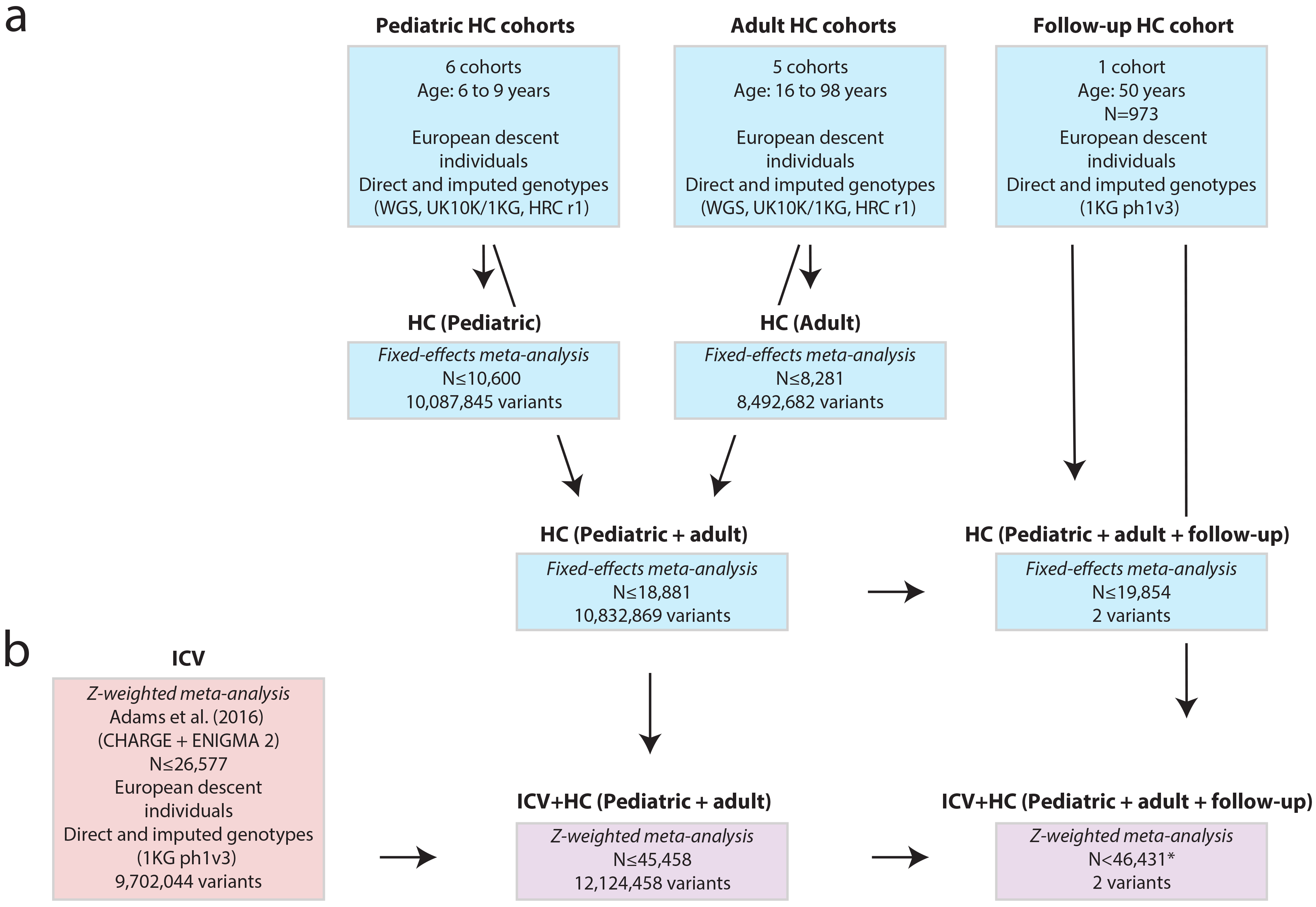
Study design. (a) Head circumference meta-analysis design using a fixed-effect metaanalysis including different developmental stages. (b) Combined head circumference and intracranial volume meta-analysis design using a Z-weighted meta-analysis. ICV - intracranial volume. WGS - Whole Genome Sequencing; UK10K/1KG - Joint UK10K/1000 Genomes imputation template; 1KG - 1000 Genomes imputation template; HRC- The Haplotype Reference Consortium r1. * Due to sample dropout only N≤43,529 were available

**Figure 2:**
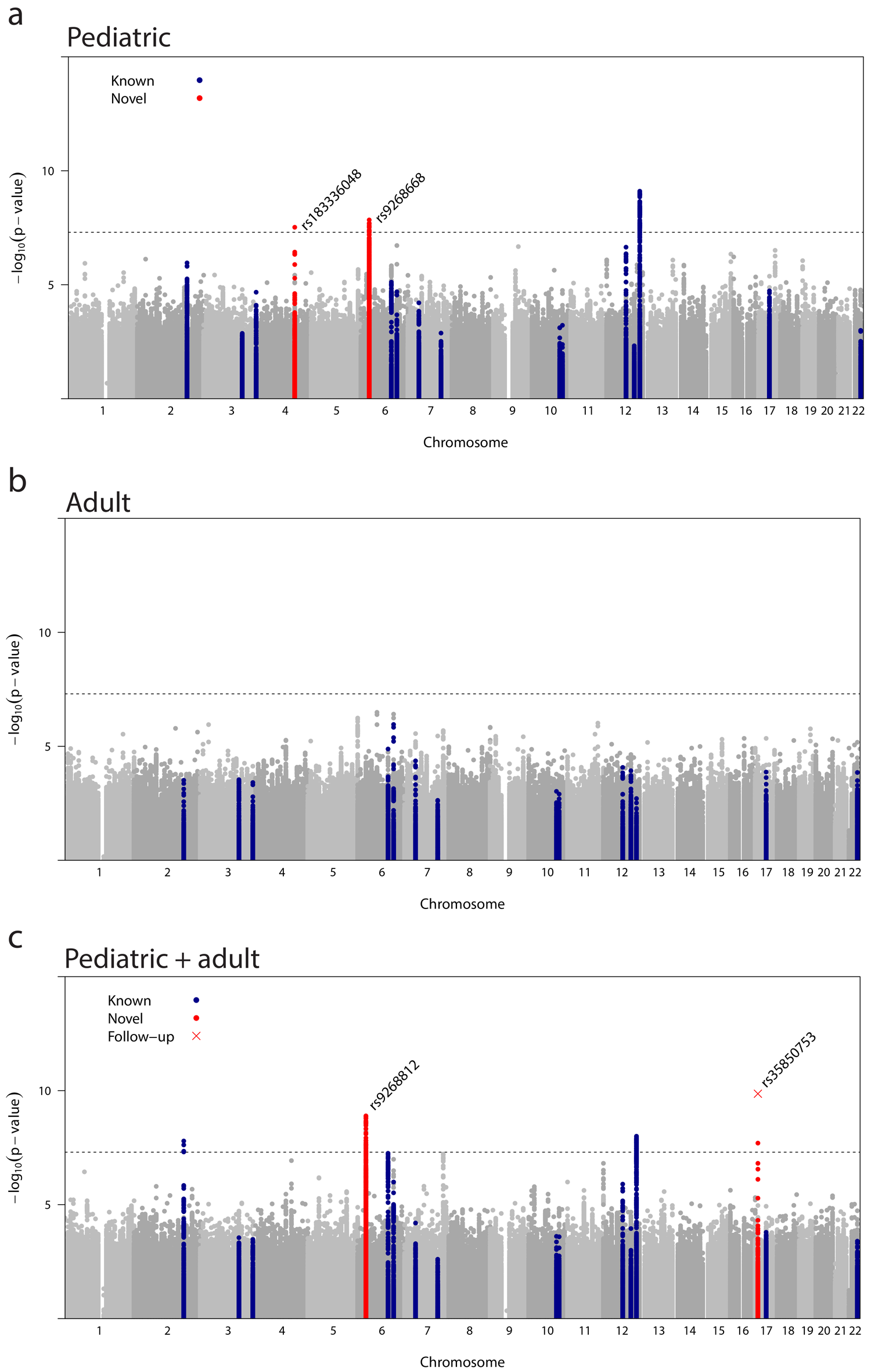
Genome-wide association with final head circumference (HC). (a) HC(Pediatric): N=8,281, (b) HC(Adult): N=10,600 and (c) HC(Pediatric+adult): N=18,881 meta-analysis. The dashed line represent the threshold for genome-wide (*p*<5.0×10^−8^) significance. Known variants for intracranial volume, brain volume and head circumference (Supplementary Table S7) are shown in blue. Novel signals (r^2^=0.2, kb=±500) passing a nominal genome-wide association threshold (*p*<5×10^−8^) are shown with their lead SNP in red. Replicated signals are labelled with a red cross. Genomic position is shown according to NCBI Build 37. Accounting for multiple testing, the adjusted level of genome-wide significance is *p*<3.3×10^−8^.

**Table 1:** Genome-wide meta-analysis of final head circumference (HC): Novel loci.

| Meta-analysis | Variant | Chr | Position (b37) | EA | EAF | Discovery |  |  |  | Follow-up |  |  | Discovery + follow-up |  |  |
| --- | --- | --- | --- | --- | --- | --- | --- | --- | --- | --- | --- | --- | --- | --- | --- |
| | | | | | | $\beta$ (SE) | $p$ | N | $p_{Het}$ | $\beta$ (SE) | $p$ | N | $\beta$ (SE) | $p$ | |
| *HC(Pediatric)* | rs183336048 | 4 | 128302747 | T | 0.02 | 0.31 (0.06) | $3.0 \times 10^{-8}$ | 10,303 | 0.62 | - | - | - | - | - | - |
| *HC(Pediatric+adult)* | rs9268812<sup>a</sup> | 6 | 32424568 | A | 0.35 | 0.07 (0.01) | $1.3 \times 10^{-9}$ | 18,881 | 0.85 | 0.01(0.05) | 0.87 | 973 | 0.07(0.01) | $2.1 \times 10^{-9}$ | 19,557 |
| *HC(Pediatric+adult)* | rs35850753 | 17 | 7578671 | T | 0.02 | 0.22 (0.04) | $2.0 \times 10^{-8}$ | 18,584 | 0.21 | 0.64(0.16) | $4.5 \times 10^{-5}$ | 973 | 0.24(0.04) | $2.1 \times 10^{-10}$ | 19,557 |

<sup>a</sup>rs9268812 is in linkage disequilibrium with rs9268668 ( $r^2 = 0.60$ ) that passes the genome-wide significance threshold within the HC(Pediatric) meta-analysis (Supplementary Table S2).
Position, base pair position; EA, effect allele; EAF, effect allele frequency; $\beta$ , effect estimate; SE standard error; $p$ , p-value; N, sample size; $p_{Het}$ , p-value for heterogeneity statistic (based on Cochran's Q-test for heterogeneity).

Growth curve modelling of HC scores between birth and the age of 15 years in participants of the ALSPAC sample, using a stratified Super Imposition by Translation And Rotation (SITAR) model [18], suggested that carriers of the T risk allele at rs35850753 developed larger heads from mid childhood onwards (Figure 4), with risk alleles being positively related to individual differences in mean HC (*p*=6.9×10^−12^) and HC growth velocity (*p*=7.1×10^−11^, Supplementary Table S6). For example, at the age of 10 years male carriers had a HC score of 54.16 cm and non-carriers a score of 53.63 cm. In comparison, female carriers and non-carriers had a score of 53.21 cm and 52.74 cm respectively. rs3585075 resides within the tumor suppressor encoding *TP53* gene and is not related to any known GWAS locus for HC, ICV, brain volume (Supplementary Table S7) nor any locus affecting height [19] when conducting a conditional analysis (data not shown). In addition to these novel associations, our analysis replicated known signals for infant HC on chromosome 12q24.31 [13] and a previously reported joint signal of infant HC and adult ICV on chromosome 2q32.1 [12] (Figure 2c).

**Figure 3.**
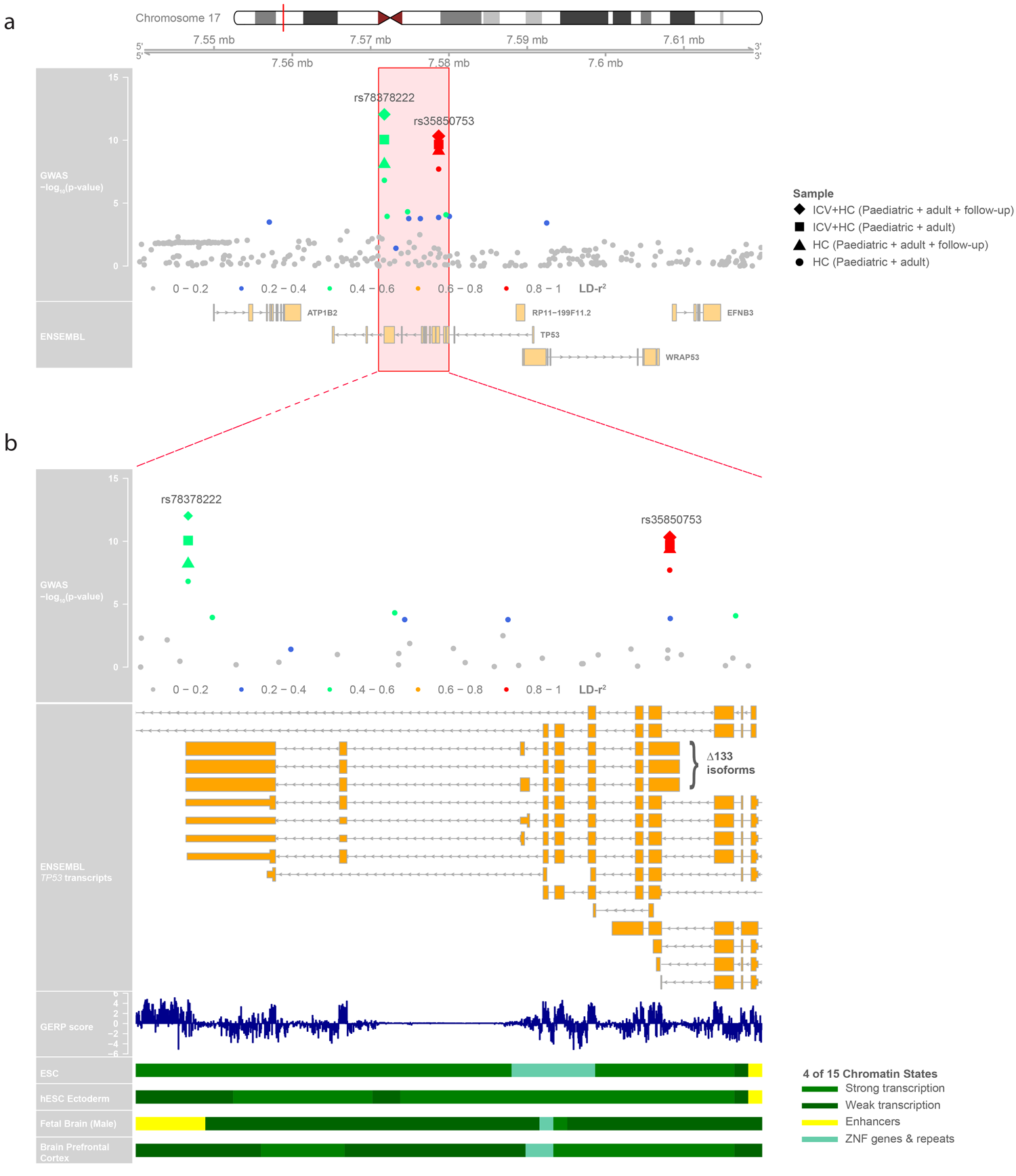
Regional association plot at 17p13.1 associated with final head circumference (HC) and final cranial dimension. (a) Depicts a 800 Mb window and (b) a zoomed view of genetic association signals and functional annotations near *TP53*. Within each plot, in the first panel SNPs are plotted with their −log_10_ p-value as a function of the genomic position (b37). This panel shows the statistical evidence for association based on the fixed-effect meta-analysis HC (Pediatric+adult) samples, but also for lead signals from including HC follow-up and intracranial volume (ICV) samples (see legend for shape coding). SNPs are colored based on their correlation with the HC lead signal (rs35850753, pairwise LD-r^2^-values). The second panel represents the gene region (ENSEMBL GRCh37). The third panel in (b) presents the Genomic Evolutionary Rate Profiling (GERP++) score of mammalian alignments. The last four panels in (b) show 4 of 15 core chromatin states, present in the zoomed view, from the Roadmap Epigenomics Consortium including Embryonic Stem Cells (ESC), hESC Derived CD56+ Ectoderm Cultured Cells, Fetal Brain (Male) and Brain Dorsolateral Prefrontal Cortex respectively (see legend for color coding).

**Figure 4.**
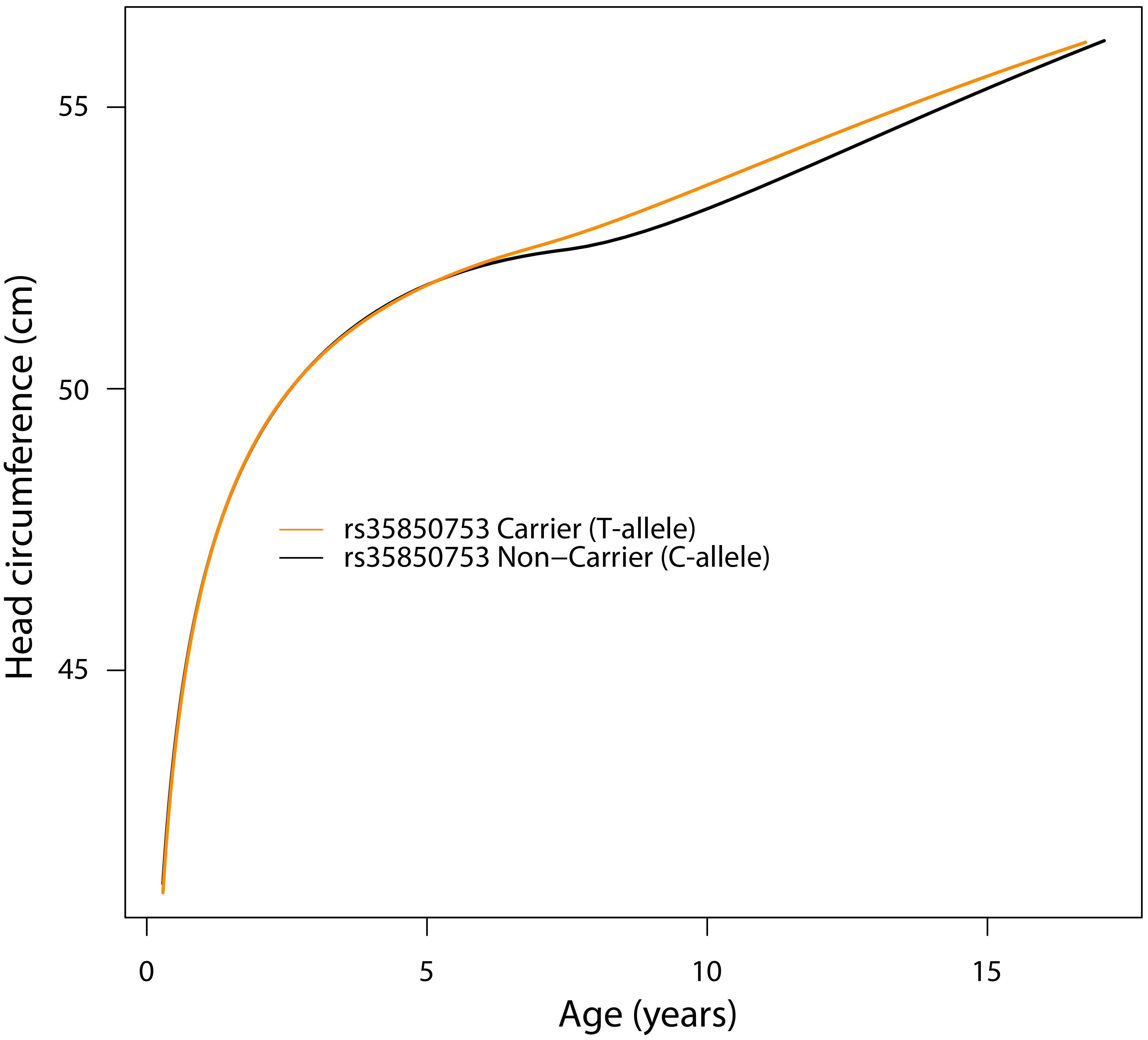
Stratified head circumference growth model trajectories for rs35850753 carriers (T allele) versus non-carriers (C-allele). The growth model was based on untransformed head circumference (cm) scores spanning birth to 15 years observed in 6,225 ALSPAC participants with up to 13 repeat measures (17,269 observations) using a mixed effect SuperImposition by Translation And Rotation (SITAR) model.

Applying a gene-based test approach [20], multiple HC-associated genes were identified (Supplementary Table S8-10). The strongest signal in HC (Pediatric), and to a lesser extent in HC (Pediatric+adult), resides at 12q24.31 (lead gene-wide signal *MPHOSPH9*, *p*=2.3×10^−10^) and contains single variants in linkage disequilibrium (LD) with known GWAS signals for infant HC [13] (e.g. *SBNO1*, *p*= 2.0×10^−7^). The strongest gene-wide signals that did not harbor variants in LD with known or novel single GWAS variants were identified at 5q31.3 (lead gene-wide signal *SLC4A9 p*=6.6×10^−9^), and at 16p 13.3 (lead gene-wide signal *E4F1 p*=1.6×10^−8^), using summary statistics from HC (Pediatric+adult) meta-analyses. Gene-based analyses were complemented with studies predicting gene expression levels in multiple tissues (Supplementary Table S11). Notably, for the HC (Pediatric) gene-wide signal at 6p21.32, including the *PRRC2A* locus (gene-wide signal *p*=7.2×10^−7^), predicted gene expression levels in whole blood were found to be inversely associated with HC scores, using S-PrediXcan [21] software (*p*= 5.7×10^−7^; Supplementary Table S11).

### Genetic architecture of HC scores during development

Linkage-disequilibrium score regression (LDSC) [22] analyses (Figure 5a, Supplementary Table S12) using genome-wide summary statistics suggested that heritability estimates during childhood (6 to 9 years) are higher (SNP-h^2^=0.31(SE=0.05)) than in adult samples (16 to 98 years; SNP-h^2^=0.097(SE=0.06)), although 95% confidence intervals marginally overlap. The estimated genetic correlation [23] between both developmental windows was high (LDSC-r_g_=1.04(SE=0.39), *p*=0.0075). The LD-score regression intercepts were consistent with one for all HC meta-analyses, suggesting little inflationary bias in GWAS (Supplementary Table S12).

To investigate developmental changes in the genetic architecture of HC scores, we carried out a multivariate analysis of genetic variances using genetic-relationship-matrix structural equation modelling (GSEM) [24]. Fitting a saturated Cholesky decomposition model (Figure 5b) to HC scores assessed in ALSPAC participants (N=7,924) at the ages of 1.5, 7 and 15 years (Supplementary Table S13), we observed total SNP-h^2^ estimates of 0.35 (SE=0.07), 0.43 (SE=0.05), and 0.39 (SE=0.07) respectively (Figure 5c). More importantly, this analysis suggested that a large proportion of genetic factors contributing to phenotypic variation in HC scores remains unchanged during the course of development, with genetic factors operating at the age of 1.5 years explaining 63.1% (SE=9%) and those at age 7 years 76.5% (SE=5%) of the genetic variance at age 15 years, respectively. Consistently, strong genetic correlations were identified among all scores during development (1.5-7 years, r_g_=0.89 (SE=0.07); 1.5-15 years, r_g_=0.79 (SE=0.09); 7-15 years, r_g_=0.87 (SE=0.04), in support of LD-score correlation analyses.

**Figure 5:**
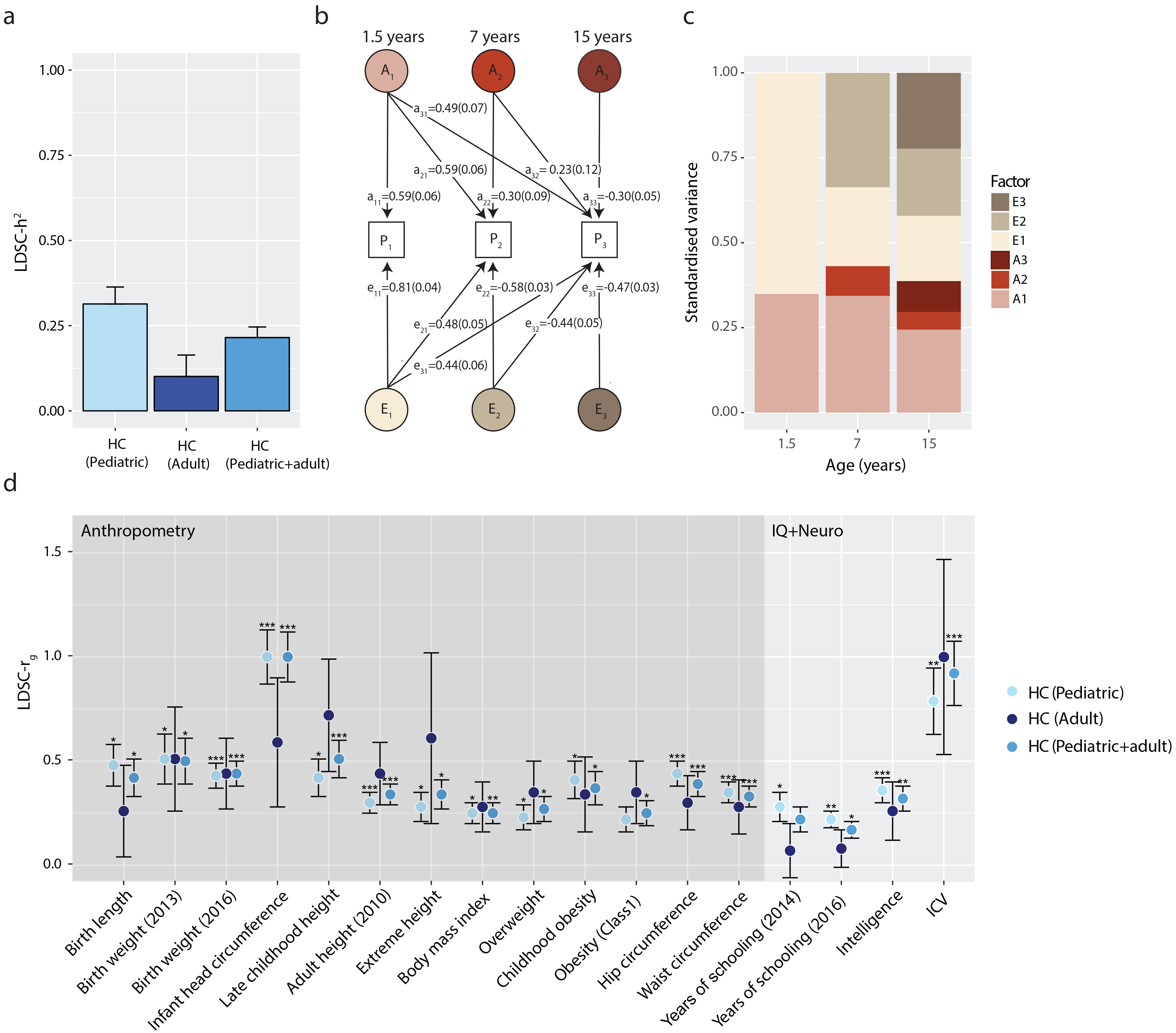
Genetic architecture of head circumference (HC). (a) Linkage-disequilibrium score SNP-heritability (LDSC-h^2^) for HC (Pediatric), HC (Adult) and HC (Pediatric+adult) meta-analyses. (b) Genetic-relationship matrix structural equation modelling (GSEM) of head circumference during development: Path diagram of the full Cholesky decomposition model using longitudinal head circumference measures from ALSPAC (1.5 years (N =3,945), 7 years (N=5,819), and 15 years (N=3,406)). Phenotypic variance (P1, P2, P3) was dissected into genetic (A1, A2 and A3) and residual (E1, E2 and E3) factors. Observed measures are represented by squares and latent factors by circles. Single headed arrows (‘paths’) define relationships between variables. The variance of latent variables is constrained to unit variance. (c) Standardized genetic and residual variance components for head circumference during development. Variance components were estimated using the GSEM model as shown in b. (d)Linkage-disequilibrium score correlation (LDSC-r_g_) for HC(Pediatric), HC(Adult) and HC(Pediatric+adult) and 235 phenotypes: 17 genetic correlation estimates passing a Bonferroni threshold (*p*<0.00014) are shown. *** *p*<10^−8^; \*\**p*<10^−5^; \**p*<0.00014

### Genetic correlation of complex phenotypes with HC

A systematic screen for genetic correlations between HC scores and 235 complex phenotypes using LD score regression [23], identified moderate to strong positive genetic correlations (r_g_≥0.3) with many anthropometric and cognitive / cognitive proxy traits. This includes HC scores during infancy, birth weight, birth length, height, extreme height, hip circumference, childhood obesity, waist circumference, intelligence scores and ICV (Supplementary Table S14, Figure 5d). Weaker positive genetic correlations (0<r_g_<0.3) were also present for years of schooling, obesity, body mass index and overweight.

The strongest cross-trait genetic correlation was identified between HC (Pediatric+adult) and ICV (r_g_=0.91(SE=0.16), *p*=1.6×10^−8^). However, there was little evidence that SNP-h^2^ estimates for HC are enriched for genes that are highly expressed in brain tissues or chromatin marks in neural and bone tissue/cell types, beyond chance (data not shown) studying summary statistics from either of the conducted HC meta-analyses.

### Combined genome-wide analysis of HC scores and ICV

Given the prior expectation of similar genetic architectures between HC and ICV, supported through genetic correlation analyses, we meta-analyzed both phenotypes by combining HC summary statistics from pediatric and adult cohorts (N=18,881) with ICV summary statistics from the CHARGE and ENIGMA2 consortia [12] (N=26,577, Figure 1b) using a Z-score weighted meta-analysis (Figure 6, Table 2, Supplementary Tables S15, S16). The strongest evidence for novel genetic association in this combined cranial dimension analysis was observed for the low-frequency marker rs78378222 (EAF=0.02; *p*=7.9×10^−11^) at the 17p 13.1 locus, a functional variant that is in LD with rs35850753, the strongest GWAS signal for HC (r^2^=0.56, *p*=3.6×10^−9^, Figure 3a and b). In addition, we identified 8 independent genetic loci not previously reported for either HC or ICV (Supplementary Table S15). Evidence for association at rs9268812, identified in the HC (Pediatric+adult) meta-analysis, decreased, although it still passed the threshold for genome-wide significance.

**Figure 6:**
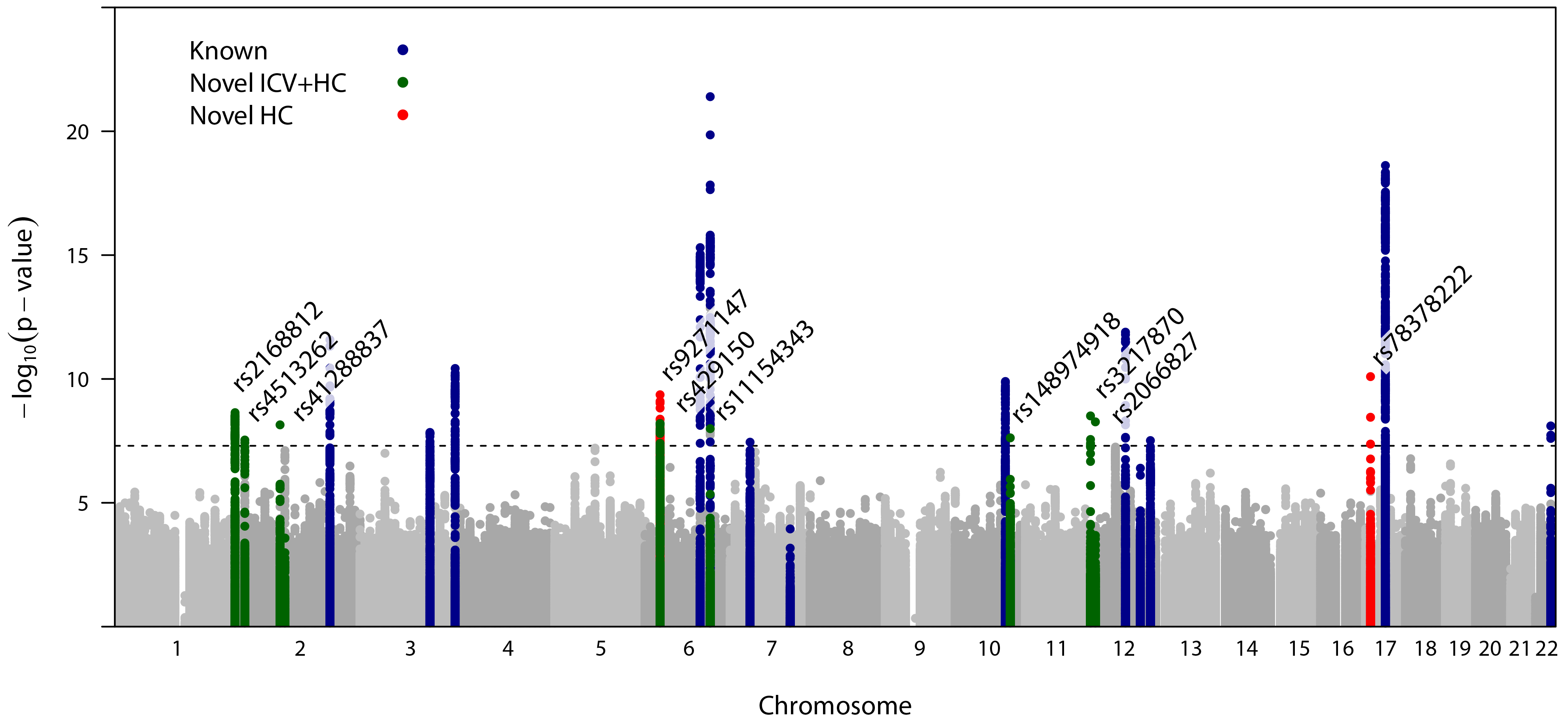
Genome-wide association analysis of final cranial dimension. A genome-wide weighted Z-score meta-analysis of combined head circumference (HC) and intracranial volume (ICV) was carried out (ICV+HC (Pediatric+adult): N= 45,458). The dashed line represents the threshold for genome-wide (*p*<5.0×10^−8^) significance. Known variants for ICV, brain volume and HC are shown in blue (Supplementary Table S7). Novel signals (r^2^=0.2, kb=±500) passing a nominal genome-wide association threshold (p<5 ×10^−8^) are shown with their lead SNP in green (Table 2, Table S15). HC (Pediatric + adult) signals identified in this study are shown in red. Genomic position is shown according to NCBI Build 37. Accounting for multiple testing, the adjusted level of genome-wide significance is *p*<3.3×10^−8^.

**Table 2:** Genome-wide combined meta-analysis of intracranial volume (ICV) and head circumference (HC): Novel loci.

| Variant | Chr | Position(b37) | EA | EAF | ICV |  |  | HC (Pediatric + adult) |  |  | ICV+HC (Pediatric + adult) |  |  |
| --- | --- | --- | --- | --- | --- | --- | --- | --- | --- | --- | --- | --- | --- |
| | | | | | $\beta^*$ (SE) | *p* | N | $\beta^*$ (SE) | *p* | N | $\beta^*$ (SE) | *p* | N |
| rs2168812 | 1 | 243777066 | A | 0.18 | 0.05(0.01) | $9.0 \times 10^{-6}$ | 26,577 | 0.06(0.01) | $1.5 \times 10^{-5}$ | 18,881 | 0.05(0.01) | $2.3 \times 10^{-9}$ | 45,458 |
| rs4513262 | 2 | 10181457 | T | 0.21 | 0.05(0.01) | $7.7 \times 10^{-7}$ | 26,455 | 0.04(0.01) | $2.3 \times 10^{-3}$ | 18,881 | 0.05(0.01) | $2.9 \times 10^{-8}$ | 45,336 |
| rs41288837 | 2 | 85531116 | T | 0.03 | -0.10(0.03) | $1.0 \times 10^{-4}$ | 26,179 | -0.15(0.03) | $1.6 \times 10^{-6}$ | 17,737 | -0.11(0.02) | $7.2 \times 10^{-9}$ | 43,916 |
| rs429150<sup>a</sup> | 6 | 32075563 | T | 0.52 | 0.04(0.01) | $3.1 \times 10^{-5}$ | 24,242 | 0.04(0.01) | $1.0 \times 10^{-4}$ | 18,881 | 0.04(0.01) | $3.9 \times 10^{-8}$ | 43,122 |
| rs9268812 | 6 | 32424568 | A | 0.35 | 0.02(0.02) | $3.0 \times 10^{-1}$ | 59,57 | 0.07(0.01) | $1.3 \times 10^{-9}$ | 18,881 | 0.05(0.01) | $1.6 \times 10^{-8}$ | 24,838 |
| rs11154343 | 6 | 126412953 | T | 0.31 | -0.05(0.01) | $1.7 \times 10^{-8}$ | 26,577 | -0.03(0.01) | $1.2 \times 10^{-2}$ | 18,881 | -0.04(0.01) | $1.0 \times 10^{-8}$ | 45,458 |
| rs7808664<sup>a</sup> | 7 | 32917550 | A | 0.37 | -0.04(0.01) | $1.2 \times 10^{-6}$ | 26,577 | -0.03(0.01) | $2.0 \times 10^{-3}$ | 18,881 | -0.04(0.01) | $3.6 \times 10^{-8}$ | 45,458 |
| rs148974918 | 10 | 112033051 | T | 0.05 | -0.08(0.02) | $9.4 \times 10^{-5}$ | 26,577 | -0.1(0.02) | $1.6 \times 10^{-5}$ | 18,881 | -0.08(0.02) | $2.4 \times 10^{-8}$ | 45,458 |
| rs3217870 | 12 | 4400111 | T | 0.62 | -0.03(0.01) | $1.0 \times 10^{-4}$ | 26,577 | -0.05(0.01) | $7.0 \times 10^{-7}$ | 18,881 | -0.04(0.01) | $3.1 \times 10^{-9}$ | 45,458 |
| rs2066827 | 12 | 12871099 | T | 0.77 | 0.05(0.01) | $3.9 \times 10^{-6}$ | 19,990 | 0.05(0.01) | $9.3 \times 10^{-5}$ | 18,881 | 0.05(0.01) | $5.4 \times 10^{-9}$ | 38,871 |
| rs35850753<sup>b</sup> | 17 | 7578671 | T | 0.02 | 0.1(0.03) | $1.5 \times 10^{-3}$ | 23,972 | 0.21(0.04) | $2.0 \times 10^{-8}$ | 18,584 | 0.14(0.02) | $3.6 \times 10^{-9}$ | 42,556 |
| rs78378222<sup>b</sup> | 17 | 7571752 | T | 0.02 | 0.15(0.04) | $1.4 \times 10^{-5}$ | 23,634 | 0.21(0.04) | $1.5 \times 10^{-7}$ | 17,737 | 0.17(0.03) | $7.9 \times 10^{-11}$ | 41,371 |

ICV - GWAS statistics as reported by the intracranial volume (ICV) meta-analysis of ENIGMA2 and CHARGE samples, $N \leq 26,577$
HC (Pediatric+adult) - Combined pediatric and adult head circumference meta-analysis, $N \leq 18,881$
a - Passes a nominal threshold for genome-wide significance only ( $p < 5 \times 10^{-8}$ )
Position, base pair position; EA, effect allele; EAF, effect allele frequency; $\beta^*$ , Standardized effect estimate; SE, standard error; *p*, p-value; N, sample size

Adding further genotype information from the HC follow-up cohort (N=973, Figure 1b), strengthened the evidence for association at both rs78378222 and rs35850753 (*p*=8.8×10^−13^ and *p*=4.9×10^−11^respectively, total N≤43,529, Figure 3a and b, Supplementary Table S15) corresponding to a change in 0. 19 (SE=0.03) and 0.16 (SE=0.02) standard deviation (SD) units respectively. However, within neuroimaging samples only, support for association at rs78378222 was low in the combined CHARGE and ENIGMA2 samples [12] (Table 2). In addition, we observed increased statistical evidence for association at 9 known markers compared to the original studies for either HC, brain volume or ICV (Supplementary Table S16). Note that genetic effects with respect to a combined final cranial dimension cannot be translated into absolute units as HC scores and ICV relate to diametric versus volumetric properties respectively.

### Biological and phenotypic characterization of signals

rs78378222 resides in the 3′ untranslated region (UTR) of *TP53* and the C effect allele leads to a change in the *TP53* polyadenylation signal that results in impaired 3’-end processing for many *TP53* mRNA, while rs35850753 resides within the 5’-UTR of the Δ133 *TP53* isoforms and otherwise intronically (Figure 3b). Species comparison showed that variation at rs78378222 is highly conserved (GERP-score=5.28), while variation at rs35850753 is not (GERP-score=−2.8) (Figure 3b) [25]. According to a core 15-state chromatin model, variation at rs78378222, but not at rs35850753, is in LD (r^2^=0.8) with an enhancer in fetal brain (Figure 3b). However, using the FUMA web tool [26], we identified evidence for LD between rs35850753cis and nearby cis eQTLs at *ATP1B2*, *TP53*, *EIF4A1*, *TNFSF12*, *TNFSF12-TNFSF13*, *TNFSF13* and *NLGN2* loci in blood (Supplementary Table S17), but not for rs78378222 (data not shown). Specifically, rs35850753 is in modest LD (maximum LD r^2^=0.22) with *TP53* eQTL explaining variation in gene level *TP53* transcript in blood, with the rare T risk allele being associated with lower full-length transcript *TP53* levels. There was little support for eQTLs or meQTLs at either rs78378222 or rs35850753 in brain, according to Brain xQTL [27], when adjusted for the number of loci tested (data not shown). There was also no evidence for association with Human Embryonic Cranial Tissue Epigenomic Data [28] passing a multiple testing threshold (data not shown).

Finally, we characterized the *TP53* association signals for final HC and combined final ICV+HC phenotypically using a phenome-wide scan in the UK Biobank, as implemented in PHESANT [29] (rs35850753: Supplementary Table S18; rs78378222: Supplementary Table S19). For rs35850753, standing height is increased by 0.012 cm (SE=0.002) and sitting height by 0.015 cm (SE=0.003) for each increase in minor effect allele. The log odds of having an inpatient primary diagnosis code for “Fracture of tooth” increase by 0.42 (SE=0.078) and the log odds of a participant answering “yes” to “ever had hysterectomy” by 0.06 (SE=0.011) per effect allele. For each increase in minor effect allele of rs78378222, standing height is increased by 0.015 cm (SE=0.002) and sitting height by 0.019 cm (SE=0.003). The log odds of a participant answering “yes” to “ever had hysterectomy” by 0.065 (SE=0.011) per effect allele. Sensitivity analysis adjusting, in addition, for 10 principal components did not change the nature of these findings (data not shown).

## Discussion

Investigating up to 46,000 individuals of European descent, this study identifies and replicates evidence for genetic association between a novel region on chromosome 17p13.1 and both final HC and ICV+HC, implicating low-frequency variants of large effect within *TP53*. We furthermore demonstrate that the genetic architecture of HC is developmentally stable and genetically correlated with ICV. This is supported by the identification of 8 further common and rare independent loci that are associated with cranial dimension as a combined HC and ICV phenotype, illustrating allele frequency spectrum of the underlying genetic architecture.

For final HC, the strongest evidence for association at 17p13.1 is observed with rs35850753, while final cranial dimension is most strongly associated with rs78378222. Both rs35850753 and rs78378222 are low-frequency variants, in partial linkage disequilibrium, and their effect sizes are substantially larger than any previously reported GWAS signals for either HC or ICV alone, reaching nearly a fifth and quarter of a SD unit change in final cranial dimension and final HC per rare effect allele respectively. For HC, this translates into an increase of approximately 0.5 cm in HC between carriers and non-carriers of rare alleles at the age of 10 years. Based on longitudinal analyses, it is most likely that genetic effects of rs35 850753 on final HC start to emerge during mid-childhood, while we have no comparable longitudinal data source available to evaluate trajectory effects on cranial dimension.

*TP53* encodes the p53 protein, a transcription factor that binds directly and specifically as a tetramer to DNA in a tissue- and cell-specific manner and has a range of anti-proliferative functions, lending it the nickname ‘guardian of the genome’. The activation of p53 in response to cellular stress promotes cell cycle arrest, DNA repair and apoptosis [30]. *TP53* mutations are present in approximately 30% of tumor samples making it one of the most studied genomic loci with over 27,000 somatic and 550 germline mutations described to date (source - IARC TP53 database [31]). The low-frequency C allele at rs78378222 leads to a change in the *TP53* polyadenylation signal that results in impaired 3’-end processing and termination of many *TP53* mRNA isoforms [32], including full length *TP53* isoforms, although rs78378222 is also in LD with an enhancer region in fetal brain. In contrast, rs35850753 resides in the 5´ UTR of *TP53Δ133* isoforms that are transcribed by an alternative promoter. This leads to the expression of an N-terminally truncated p53 protein, initiated at codon 133, lacking the trans activation domain [33]. *TP53 Δ133* isoforms are known to directly and indirectly modulate p53 activity and differentially regulate cell proliferation, replicative cellular senescence, cell cycle arrest and apoptosis in response to stress such as DNA damage, including the inhibition of tumor suppressive functions of full-length p53 [34]. This mechanism is consistent with the observed link between the rs35850753 low-frequency T allele and lower full length *TP53* transcript level. Thus, rare effect alleles at both rs78378222 and rs35850753 could potentially, via different biological mechanisms, be linked to impaired p53 activity and thus heightened proliferative potential and less apoptosis of normal human cells, consistent with larger HC scores and a larger cranial dimension.

It is noteworthy that rs78378222 has been associated in particular to risk of cancers of the nervous system including glioma [35] and both, rs35850753 and rs78378222 have been robustly associated with neuroblastoma [36]. Evidence for neurological phenotypic consequences of *TP53* variation has recently been strengthened by the discovery of *TP53* as a risk locus for general cognitive function using a gene-based approach [37]. *TP53* knockout mouse embryos show furthermore broad cranial defects involving skeletal, neural and muscle tissues [38]. Similarly, mouse models for Treacher Collins syndrome (a disorder of cranial morphology which arises during early embryological development as a result of defects in the formation and proliferation of neural crest cells) could be rescued by inhibition of p53 during embryological patterning [39]. In particular, there is support from animal and tissue models for a role of p53 in neural crest cell (NCC) development [38] with NCCs supplementing head mesenchyme during fetal development [11, 40]. NCCs also contribute to the development of a thick 3-membrane layer called the meninges, that cover the telencephalon [11, 40] and directly locate underneath the skull. During postnatal brain growth, the calvarial bones are drawn outward, partially due to the expanding meninges, triggering the production of membranous skull bone [40]. Moreover, meninges have been thought to play a key role in the coordinated integration of signaling pathways regulating both neural and skeletal cranial growth [11]. It is possible to speculate that p53 is part of these joint regulatory mechanisms, for example, via Wnt signaling regulation [41]. However, beneficial effects of rs35850753 and rs78378222 on growth patterning leading to an increased HC and cranial dimension might be counterbalanced by adverse outcomes such as glioblastoma, keeping both variants at a lower frequency.

Combined analysis of HC and ICV, as related measures of final cranial dimension, also identified association at 8 further loci, in addition to variation at 17p13.1 and loci previously reported for either infant HC and/or adult ICV. This includes the low-frequency variant rs41288837, exerting moderately large effects that correspond to an approximately 10% decrease in SD units of final cranial dimension per rare T allele. rs41288837 is a missense variant in *TCF7L1* at 2p11.2, a locus encoding a transcription factor mediating Wnt signaling pathways that are known to play an important role in vertebrate neural development [42]. The effects of this variant were consistent for both HC and ICV, although each of the individual trait analyses was too underpowered to detect association at this variant at a genome-wide level. An additional low-frequency variant in this study, rs183336048 at 4q28.1, was identified as associated with pediatric HC only, but could not be replicated due to a lack of comparable age-matched follow-up cohorts. rs183336048 lies 5’ to *INTU* which encodes a polarity effector ‘inturned’ with roles in neural tube patterning and cilliation [43]. Common variants identified in the ICV+HC combined metaanalysis also include intronic variation in *AKT3* (rs2168812) and *CCND2* (rs3217870), which are related through the phosphatidylinositol 3-kinase (PI3K-AKT) pathway [44]. Disruption of PI3K-AKT pathway components causes megalencephaly-polymicrogyria-polydactyly-hydrocephalus syndrome and a spectrum of related megalencephaly syndromes [45, 46]. This supports previous *in silico* pathway analysis, which nominated PI3K-AKT as candidate pathway for intracranial volume [12].

Genetic correlation analyses provided strong evidence for shared genetic determinants between HC and both anthropometric (birth weight, height, waist and hip circumference) and cognitive traits, as well as ICV. Genetic correlations with waist circumference and hip circumference recapitulate observed correlations between the size of the maternal pelvis and the size of the neonatal cranium [47], possibly induced because bipedal locomotion limits pelvic size. We also confirmed previously-reported genetic links between educational attainment and infant HC [48], and identified additional evidence for genetic correlation between HC and intelligence, especially pediatric HC. With strong shared genetic liability (genetic correlation coefficient near one), we considered HC and ICV to be related proxy measures of an underlying phenotype, which we termed final cranial dimension. This also suggests that estimated skeletal volume, a combination of HC, cranial height and cranial length [49], might represent a more accurate, easily accessible and inexpensive measure to enhance power for future genetic analysis using a multi-trait approach in combination with ICV [50], exploiting similar volumetric properties.

Multivariate analyses of genetic variance showed that genetic factors contributing to variation in HC during infancy explain the majority of genetic variance during later life, although novel genetic influences arise both during mid-childhood and adolescence. This is further reflected in strong GSEM-based genetic correlations across childhood and adolescence, and strong LDSC-based genetic correlations between infant and adult HC. The estimated LDSC-heritability of HC in adult samples was lower than in pediatric samples, with only marginally overlapping 95% confidence intervals, implying that phenotypic variation in final HC is less well accounted for by genetic influences than in childhood, probably as skeletal growth processes have ceased.

The discovery that low-frequency variation, especially near *TP53*, is associated with HC demonstrates the scientific value of testing for variation in the lower allele frequency spectrum and the utility of comprehensive imputation templates. Low-frequency variants identified in this study had larger effects than common variants (Supplementary Figures S7 and S8), in keeping with findings from a range of complex phenotypes including anthropometric traits [17, 51, 52]. Nevertheless, despite having sufficient power to detect low-frequency variation explaining as little as 0.11% of the variance in HC, this study was underpowered for rare variant analysis (Supplementary Note 4), underlining the need for even larger research efforts. Collectively, our findings provide novel insight into the genetic architecture of cranial development and contribute to an improved understanding of its dynamic nature throughout human growth and development.

## Methods

### Study population

For the discovery analysis, we adopted a two-stage developmental design including cohorts with HC scores during childhood (Pediatric HC, mean age 6 to 9 years of age, N=10,600), during adulthood (Adult HC, mean age 44 to 61 years of age, N=8,281) and a combination thereof (N=18,881) including individuals of European descent from 11 population-based cohorts (Supplementary Table S1, S5, Figure 1). Cohorts include The Avon Longitudinal Study of Parents and Children (ALSPAC), the Generation R Study (GenR), the Western Australia Pregnancy Cohort Study (RAINE), the Copenhagen Prospective Study on Asthma in Children (COPSAC2000 and COPSAC2010), the Infancia y Medio Ambiente cohort (INMA), the Hellenic Isolated Cohorts HELIC-Pomak and HELIC-MANOLIS, the Orkney Complex Disease Study (ORCADES), the Croatian Biobank Korčula (CROATIA-KORCULA), and the Viking Health Study-Shetland (VIKING). Within ALSPAC analysis was performed separately in individuals with whole-genome sequence data (ALSPAC WGS) and chip-based genotyping (ALSPAC GWA). For follow-up, we studied 973 individuals from the Croatian Biobank, Split (CROATIA-SPLIT) (Supplementary Table S5). Institutional and/or local ethics committee approval was obtained for each study. Written informed consent was received from every participant within each cohort. An overview of each cohort can be found in Supplementary Table S1 and S5 with more detailed information in the Supplementary Note 1.

### Genotyping

Within ALSPAC WGS, whole-genome sequencing data was obtained using a low read depth (average 7x) strategy as previously described [52]. Chip based genotyping was performed on various commercial genotyping platforms, depending on the cohort (Supplementary Table S1). Prior to the imputation, all cohorts had similar quality control; variants were excluded because of high levels of missingness (SNP call rate <98%), strong departures from Hardy-Weinberg equilibrium (*p*<10^−6^), or low MAF (<1%). Individuals were removed if there were sex discordance, high heterozygosity, low call rate (<97.5%) or duplicates. For imputation, the reference panel was either joint UK10K/1000 Genomes [52] or the Haplotype Reference Consortium [15]. Additional details can be found in Supplementary Table S1 and Supplementary Note 1.

In addition to study-specific quality control measures, central quality control was performed using the EasyQC R package [53]. First, variants were filtered for imputation quality score (imputed studies only, INF0>0.6), minor allele count (MAC; ALSPAC WGS MAC>4, all imputed studies MAC>10) and a minimum MAF of 0.0025. SNPs with MAF discrepancies (>0.30) compared to the HRC panel were also excluded. Marker names were harmonized and reported effect and non-effect alleles were compared against reference data (Build 37). Variants with missing or mismatched alleles were dropped, in addition all insertion/deletions (INDELs), duplicate SNPs and multiallelic SNPs were excluded. The reported effect allele frequency for each study was plotted against the frequency in the HRC reference data to identify possible strand alignment issues (Supplementary Figure S1, S3). The final number of variants passing all quality control tests and the per-study genomic inflation factor (λ) are reported in Supplementary Table S1 and S5.

### Phenotype preparation

Pertinent to this study, HC measures in all individual cohorts were transformed into Z-scores using a unified protocol. After the removal of outliers (+/− 4 SD within each sample), HC was adjusted for age within males and females separately. Residuals for each sex were subsequently transformed into Z-scores and eventually combined (thus removing inherent sex-specific effects). Note that the phenotype transformation within ALSPAC was jointly carried out for both sequenced and genome-wide imputed samples.

### Genetic-relationship structural equation modelling

Developmental changes in the genetic architecture of HC scores between the ages of 1.5 and 15 years were modelled using genetic-relationship structural equation modelling (GSEM) [24]. This multivariate analysis of genetic variance combines whole-genome genotyping information with structural equation modelling techniques using a full information maximum likelihood approach [24]. Changes in genetic variance composition were assessed with longitudinal HC scores in ALSPAC participants (7924 individuals with up to three measures; 1.5 years, N=3,945; 7 years N=5,819; 15 years, N=3,406). HC scores were Z-standardised at each age, as described above. Genetic-relationship matrices were constructed based on directly genotyped variants in unrelated individuals, using GCTA software [54], and the phenotypic variance dissected into genetic and residual influences using a full Cholesky decomposition model [24].

### Multiple testing correction

Using Matrix Spectral Decomposition (matSpD) [55], we estimated that we analyzed 1.52 effective independent phenotypes within this study (Pediatric, Adult and Pediatric+adult HC scores and ICV [12] scores) according to LDSC-based genetic correlations [22].

### Single variant association analysis

Single variant genome-wide association analysis, assuming an additive genetic model, was carried out independently within each cohort using standard software (Supplementary Table S1, Supplementary Note 1). Residualized HC scores (Z-scores) were regressed on genotype dosage using a linear regression framework. For cohorts with unrelated subjects (Supplementary Table S1) association analysis was carried out using SNPTEST v2.5.0 (-method expected, -frequentist) [56]. Note that HC scores in GenR were, in addition, adjusted for four principal components. Cohorts with related participants (HELIC cohorts) utilized a linear mixed model to control for family and cryptic relatedness, implemented in GEMMA [57].

Individual cohort level summary statistics for HC were combined genome-wide with standard error-weighted fixed effects meta-analysis, allowing for the existence of age-specific effects through an age-stratified design (Figure 1a). We restricted each HC meta-analysis (Pediatric, Adult, Pediatric+adult) to variants with a minimum sample size of N>5000. Genomic control correction was applied at the individual cohort level and heterogeneity between effects estimates was quantified using the I-squared statistic as implemented in METAL [58]. Accounting for the effective number of independent phenotypes studied, the threshold for genome-wide significance was fixed at 3.3×10^−8^ and the threshold for suggestive evidence at 6.6×10^−6^.

We contacted all studies (known to us) with a) HC information available in later childhood or adult samples b) participants of European ancestry and c) genotype data. Studies with whole-genome sequencing or densely imputed genotype data (HRC or UK10K/1KG combined templates) were included in the HC meta-analysis, while studies with imputation to other templates were reserved for follow-up. Following this strategy, the majority of studies were included in meta-analysis, with follow-up in a single study.

### Identification of known variants and conditional analysis

Known GWAS signals (*p*≤5×10^−8^) were identified from previous studies on HC in infancy [13], ICV [12, 59–61] and brain volume [62] using either published or publicly available data (Supplementary Table S7). Conditional analysis was performed with GCTA software using summary statistics from HC (Pediatric) and HC (Pediatric+adult) meta-analyses (Supplementary Figure S4). In addition, we carried out an LD clustering of independent signals from the HC (Pediatric+adult) meta-analysis with respect to all known loci. Briefly, LD clustering is an iterative process that starts with the most significant SNP, which is clumped with variants that have pairwise LD of r^2^ ≥ 0.2 within 500 kb using PLINK v1.90b3w, and all variants in LD are removed. Then, the same clumping procedure is repeated for the next top SNP and the iteration continues until there are no more top variants with *p*<10^−4^. For details, see Supplementary Note 2. For sensitivity analysis, we repeated the LD clustering with known loci for height as identified through the GIANT consortium (697 known independent height GWAS signals [19], r^2^=0.2, +/−500 kb).

### Combined meta-analysis of head-circumference and intracranial volume

We carried out a weighted Z-score meta-analysis of the combined HC (Pediatric+adult) metaanalysis and the largest publicly available genome-wide statistics on intracranial volume (ICV; N=26,577) based on data from Cohorts for Heart and Aging Research in Genomic Epidemiology (CHARGE) and the Enhancing NeuroImaging Genetics through Meta-Analysis (ENIGMA) consortium [12]. A weighted Z-score meta-analysis was carried out using METAL [58] using standardized regression coefficients and 12,124,458 imputed or genotyped variants, assuming a genome-wide threshold of significance at *p*≤3.3×10^−8^.

We used the Z-scores (Z) from the METAL output to calculate the standardized regression coefficient (β) for each SNP and trait [63]

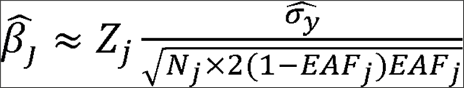

where SNP j has an effect allele frequency (EAF_j_) and 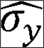 is standard deviation of the phenotype, which is assumed to equal one for standardised traits. The standard error (SE) is calculated as

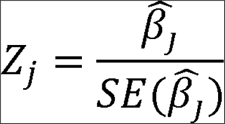

### Gene based analysis

Gene based tests for association were performed using MAGMA [20], which calculates gene-based test statistics from SNP-based test statistics, position-based gene annotations and a linkage disequilibrium reference panel of UK10K haplotypes using an adaptive permutation procedure. SNP-based test statistics were annotated using mapping files with a 50 kb symmetrical window around genes. For gene definition, we used all 19,151 protein-coding gene annotations from NCBI 37.3 and corrected for the number of genes and effective phenotypes tested, using an adjusted Bonferroni-threshold of 1.7×10^−6^.

### S-PrediXcan

We used S-PrediXcan [21] as a summary-statistic based implementation of PrediXcan to test for association between tissue-specific imputed gene expression levels and HC. This approach first predicts the transcriptome level using publicly available transcriptome datasets. Then, it infers the association between gene and phenotype of interest, by using the SNP-based prediction of gene expression as weights (predicted from the previous step) and combines it with evidence for SNP association based on phenotype-specific GWAS summary statistics. We predicted gene expression levels for cerebellum (4778 genes; GTEx v6p; Supplementary Note 3), cortex (3177 genes; GTEx v6p; Supplementary Note 3) and whole blood (6669 genes; DGN; Supplementary Note 3) using an adjusted Bonferroni threshold of *p*< 2.3×10^−6^ across all tissues tested.

### Linkage-disequilibrium score heritability and genetic correlation

Linkage-disequilibrium score regression (LDSC) [22] was carried out to estimate the joint contribution of genetic variants as tagged by common variants (SNP-h^2^) to phenotypic variation in HC. The method is based on GWAS statistics and exploits LD patterns in the genome and can distinguish confounding from polygenic influences [22]. To estimate LDSC-h^2^, genome-wide χ^2^-statistics are regressed on the extent of genetic variation tagged by each SNP (LD-score). The intercept of this regression minus one estimates the contribution of confounding bias to inflation in the mean χ^2^-statistic. LD score regression was performed with LDSC software and based on the set of well-imputed HapMap3 SNPs (~1,145,000 SNPs with minor allele frequency (MAF) > 5% and high imputation quality such as an INFO score of 0.9 or higher) and a European reference panel of LD-scores. *LD-score correlation* analysis can be used to estimate the genetic correlation (r_g_) between distinct samples by regressing the product of test statistics against the same LD-score [23]. Bivariate LD score correlation was performed with the LDHub platform [64] v1.9.0. We assessed the genetic correlation between HC scores and a series of 235 phenotypes (excluding UK Biobank) comprising anthropometric, cognitive, structural neuroimaging and other traits as described in Zheng et al. [64], with an adjusted Bonferroni threshold of *p*<1.4×10^−4^.

### Stratified LD Regression: Cell Type Expression and Epigenomics

Stratified LD score regression [65] is a method for partitioning heritability from GWAS summary statistics with respect to genes that are expressed in specific tissue/cell types. We applied this method to HC summary statistics to evaluate whether the heritability of HC is enriched for genes that are highly expressed in brain tissues. GTEx v6p (Supplementary Note 3) provided gene expression data from 13 brain tissue/cell types. Each of these tissue annotations was added to the baseline model and enrichment was calculated with respect to 53 functional categories. This is for each functional category the proportion of SNP-h^2^ divided by the proportion of SNPs in that category. We performed stratified LD score regression with independent data from the Roadmap Epigenomics consortium and ENCODE project (Supplementary Note 3), where we restricted the analysis to 55 chromatin marks identified in neural and bone tissue/cell types. Similar to the deriving enrichment in gene expression, each annotation was added to the baseline model. Chromatin analysis includes the union and the average of cell-type specific annotations within each mark. The joint gene expression and chromatin enrichment analysis, we applied a multiple testing of *p*<4.8×10^−4^ accounting for 68 neural and bone tissues/cell types tested (data from GTEx v6p, ENCODE and Roadmap; Supplementary Note 3).

### Functional annotation for novel signals

We investigated whether the novel GWAS significant signals from both HC (Pediatric) and HC (Pediatric+adult) meta-analyses were associated with Human Embryonic Craniofacial Tissue Epigenomic Data [28]. In addition, functional consequences of novel variants were explored using two web-based tools: Brain xQTL [27] and FUMA [26]. The threshold for multiple testing for eQTL was adjusted according to the number of genes near the studied novel signals and their proxy SNPs (r^2^=0.2 and ±500kb). In analysis of Brain xQTL, we corrected for multiple testing of *p*<7.4×10^−4^ to account for 68 genes tested. For FUMA, the eQTL analysis, multiple testing of *p*<7.6×10^−4^ was applied to adjust for 42 genes and, 24 blood and brain tissues/cell types (Supplementary Note 3).

### UK Biobank phenome scan

To characterize the phenotypic spectrum of identified HC signals, we conducted a phenome scan on 2,143 phenotypes in the UK Biobank cohort [66], using PHESANT [29] software. Analyses were restricted to participants of UK ancestry (UK Biobank specified variable). One from each pair of related individuals, individuals with high missingness, heterozygosity, gender mismatch and putative aneuploidies were excluded. Genotype dosage at lead single variants identified in GWAS was converted into best-guess genotypes using PLINK v1.90b3w. Linear, ordinal logistic, multinomial logistic and logistic regressions were fitted to test the association between genotype and continuous, ordered categorical, unordered categorical and binary outcomes respectively. Analyses were adjusted for age, sex and genotyping chip, and, for sensitivity analysis, 10 principal components. A conservative Bonferroni threshold was applied accounting for a total of 11,056 tests performed and 2 genotypes tested (*p*<2.26×10^−6^).

### HC growth curve modelling

Trajectories of untransformed HC (cm) spanning birth to 15 years were modelled in 6,225 ALSPAC participants with up to 13 repeat measures (17,269 observations) using a mixed effect SITAR model [18]. SITAR comprises a shape invariant mixed model with a single fitted curve, where individual curves are matched to the mean curve by modelling differences in mean HC, differences in timing of the pubertal growth spurt and differences in growth velocity [18]. Individuals with large measurement errors, i. e. with HC scores at younger ages exceeding scores at later ages (by more than 0.5 SD of the grand mean) as well as outliers (with residuals outside the 99.9% confidence interval) were excluded. The best fitting model was identified using likelihood ratio tests and the Bayesian Information Criterion and included four fixed effects for splines, a fixed effect for differences in mean HC and a fixed effect for sex, in addition to two random effects for differences in mean HC and growth velocity. Stratified models were fitted for carriers and non-carriers of increaser-alleles at candidate loci. To examine the relationship between genotype dosage and differences in HC and growth velocity, these random effects were regressed on genotype dosage using a linear model.

### Data availability

Genome-wide summary statistics have been deposited at The Language Archive, a public data archive hosted by the Max Planck Institute for Psycholinguistics. Data are accessible with a persistent identifier: https://hdl.handle.net/1839/181fe1c9-70b8-4ede-845a-6ed0b8070f75

## Acknowledgements

We gratefully acknowledge the contributions of each of the participants, midwives, nurses, general practitioners, interviewers, computer and laboratory technicians, clerical workers, research staff for their time and efforts to make each of these studies possible.

## Avon Longitudinal Study of Parents and Children (ALSPAC)

We are extremely grateful to all the families who took part in this study, the midwives for their help in recruiting them, and the whole ALSPAC team, which includes interviewers, computer and laboratory technicians, clerical workers, research scientists, volunteers, managers, receptionists and nurses. The UK Medical Research Council and Wellcome (Grant ref: 102215/2/13/2) and the University of Bristol provide core support for ALSPAC. This publication is the work of the authors who will serve as guarantors for the contents of this paper. A comprehensive list of grants funding is available on the ALSPAC website at (http://www.bristol.ac.uk/alspac/external/documents/grant-acknowledgements.pdf). Collection of head circumference data at age 15 years was supported by Wellcome and MRC (grant 076467/Z/05/Z to G.D.S). GWAS genotype data was generated by Sample Logistics and Genotyping Facilities at Wellcome Sanger Institute and LabCorp (Laboratory Corporation of America) using support from 23andMe. This study makes use of data generated by the cohorts arm of the UK10K consortium, which was supported by core Wellcome funding for UK10K (grant WT091310).

## Generation R Study (GenR)

The Generation R Study is conducted by the Erasmus Medical Center in close collaboration with the School of Law and Faculty of Social Sciences of the Erasmus University Rotterdam, the Municipal Health Service Rotterdam area, Rotterdam, the Rotterdam Homecare Foundation, Rotterdam and the Stichting Trombosedienst & Artsenlaboratorium Rijnmond [STAR-MDC], Rotterdam. We gratefully acknowledge the contribution of children and parents, general practitioners, hospitals, midwives and pharmacies in Rotterdam. The generation and management of GWAS genotype data for the Generation R Study was done at the Genetic Laboratory of the Department of Internal Medicine, Erasmus MC, the Netherlands. We would like to thank Karol Estrada, Dr. Tobias A. Knoch, Anis Abuseiris, Luc V. de Zeeuw, and Rob de Graaf, for their help in creating GRIMP, BigGRID, MediGRID, and Services@MediGRID/D-Grid, [funded by the German Bundesministerium fuer Forschung und Technology; grants 01 AK 803 A-H, 01 IG 07015 G] for access to their grid computing resources. We thank Pascal Arp, Mila Jhamai, Marijn Verkerk, Lizbeth Herrera and Marjolein Peters for their help in creating, managing and QC of the GWAS database. We thank Jie Huang at the Wellcome Trust’s Sanger Institute, at Hinxton, U.K. for the creation of imputed data, with the support of Marijn Verkerk, Carolina Medina-Gomez, PhD, and Anis Abuseiris and their input for the analysis setup. The general design of Generation R Study is made possible by financial support from the Erasmus Medical Center, Rotterdam, the Erasmus University Rotterdam, the Netherlands Organization for Health Research and Development (ZonMw), the Netherlands Organisation for Scientific Research (NWO), the Ministry of Health, Welfare and Sport and the Ministry of Youth and Families. Additionally, the Netherlands Organization for Health Research and Development supported authors of this manuscript (ZonMw 907.00303, ZonMw 916.10159, ZonMw VIDI 016.136.361 to V.W.J., and ZonMw VIDI 016.136.367 to F.R.). V.W.J. received a Consolidator Grant from the European Research Council (ERC-2014-CoG-648916). This project also received funding from the European Union’s Horizon 2020 research and innovation programme under grant agreements No 633595 (DynaHEALTH) and No 733206 (LIFECYCLE). This publication is the work of authors and hence serve as guarantors for the information of this study in this paper.

## Western Australian Pregnancy Cohort study (Raine)

This study was supported by the National Health and Medical Research Council of Australia [grant numbers 572613, 403981 and 003209] and the Canadian Institutes of Health Research [grant number MOP-82893]. The authors are grateful to the Raine Study participants and their families, and to the Raine Study research staff for cohort coordination and data collection. The authors gratefully acknowledge the NH&MRC for their long term contribution to funding the study over the last 29 years and also the following Institutions for providing funding for Core Management of the Raine Study: The University of Western Australia (UWA), Curtin University, Raine Medical Research Foundation, UWA Faculty of Medicine, Dentistry and Health Sciences, The Telethon Institute for Child Health ResearchKids Institute, and Women and Infants Research Foundation (King Edward Memorial Hospital), Murdoch University, The University of Notre Dame (Australia), and Edith Cowan University. The authors gratefully acknowledge the assistance of the Western Australian DNA Bank (National Health and Medical Research Council of Australia National Enabling Facility).

This work was supported by resources provided by the Pawsey Supercomputing Centre with funding from the Australian Government and the Government of Western Australia. This publication is the work of authors, and hence serve as guarantors for the information of this study in this paper.

## Copenhagen Prospective Study on Asthma in Children (COPSAC:)(COPSAC2000 and COPSAC2010)

We express our deepest gratitude to the children and families of the COPSAC2000 and COPSAC2010 cohort studies for all their support and commitment. We acknowledge and appreciate the unique efforts of the COPSAC research team. All funding received by COPSAC is listed on www.copsac.com. The Lundbeck Foundation (Grant no R16-A1694); The Ministry of Health (Grant no 903516); Danish Council for Strategic Research (Grant no 0603-00280B) and The Capital Region Research Foundation have provided core support to the COPSAC research center. This publication is the work of authors, and hence serve as guarantors for the information of this study in this paper.

## Infancia y Medio Ambiente (INMA)

ISGlobal is a member of the CERCA Programme, Generalitat de Catalunya. This study was funded by grants from Instituto de Salud Carlos III (Red INMA G03/176; CB06/02/0041; PI041436; PI081151 incl. FEDER funds; PS09/00432; PI12/01890 incl. FEDER funds; CP13/00054 incl. FEDER funds, MS13/00054), CIBERESP, Generalitat de Catalunya-CIRIT 1999SGR 00241, Generalitat de Catalunya-AGAUR (2009 SGR 501, 2014 SGR 822), Fundació La marató de TV3 (090430), Spanish Ministry of Economy and Competitiveness (SAF2012-32991 incl. FEDER funds), Agence Nationale de Securite Sanitaire de l’Alimentation de l'Environnement et du Travail (1262C0010), EU Commission (261357, 308333 and 603794). This publication is the work of authors, and hence serve as guarantors for the information of this study in this paper.

## Hellenic Isolated Cohorts (HELIC)

This work was funded by the Wellcome Trust (098051) and the European Research Council (ERC-2011-StG 280559-SEPI). The MANOLIS cohort is named in honour of Manolis Giannakakis, 1978-2010. We thank the residents of the Mylopotamos villages, and of the Pomak villages, for taking part. The HELIC study has been supported by many individuals who have contributed to sample collection (including A. Athanasiadis, O. Balafouti, C. Batzaki, G. Daskalakis, E. Emmanouil, C. Giannakaki, M. Giannakopoulou, A. Kaparou, V. Kariakli, S. Koinaki, D. Kokori, M. Konidari, H. Koundouraki, D. Koutoukidis, V. Mamakou, E. Mamalaki, E. Mpamiaki, M. Tsoukana, D. Tzakou, K. Vosdogianni, N. Xenaki, E. Zengini), data entry (T. Antonos, D. Papagrigoriou, B. Spiliopoulou), sample logistics (S. Edkins, E. Gray), genotyping (R. Andrews, H. Blackburn, D. Simpkin, S. Whitehead), research administration (A. Kolb-Kokocinski, S. Smee, D. Walker) and informatics (M. Pollard, J. Randall). This publication is the work of authors, and hence serve as guarantors for the information of this study in this paper.

## Orkney Complex Disease Study (ORCADES)

The Orkney Complex Disease Study (ORCADES) was supported by the Chief Scientist Office of the Scottish Government (CZB/4/276, CZB/4/710), a Royal Society URF to J.F.W., the MRC Human Genetics Unit quinquennial programme “QTL in Health and Disease”, Arthritis Research UK and the European Union framework program 6 EUROSPAN project (contract no. LSHG-CT-2006-018947). DNA extractions were performed at the Wellcome Trust Clinical Research Facility in Edinburgh. We would like to acknowledge the invaluable contributions of the research nurses in Orkney, the administrative team in Edinburgh, the people of Orkney, and the data analysts in particular Dr Thibaud Boutin for the genotype imputation to the HRC reference panel and Dr Peter Joshi for setting up the GWAS analysis pipeline used in this analysis.

## 10001 Dalmations: The Croatian Biobank (CROATIA)

The CROATIA studies were funded by grants from the Medical Research Council (UK), from the Republic of Croatia Ministry of Science, Education and Sports (108-1080315-0302; 216-1080315-0302) and the Croatian Science Foundation (8875); and the CROATIA-Korčula genotyping was funded by the European Union framework program 6 project EUROSPAN (LSHGCT2006018947). We would like to acknowledge the invaluable contribution of Dr Ozren Polasek who led and managed the recruitment in both Korcula and Split, the recruitment team from the Croatian Centre for Global Health, University of Split for the data collection and all the participants. We are grateful to Peter Lichner and the Helmholtz Zentrum Munchen (Munich, Germany), AROS Applied Biotechnology, Aarhus, Denmark and the Edinburgh Clinical research facility (Edinburgh, United Kingdom) for SNP array genotyping. Genetic analyses were supported by the MRC HGU “QTL in Health and Disease” core programme. We acknowledge the contribution of Dr Thibaud Boutin for the HRC imputation of the CROATIA-Korcula cohort and establishment of the GWAS analysis pipeline, as well as Holly Trochet for performing genotype quality controls and imputation to the 1000G reference panel for the CROATIA-Split dataset.

## The Viking Health Study – Shetland (VIKING)

The Viking Health Study – Shetland (VIKING) was supported by the MRC Human Genetics Unit quinquennial programme grant “QTL in Health and Disease”. DNA extractions and genotyping were performed at the Edinburgh Clinical Research Facility, University of Edinburgh. We would like to acknowledge the invaluable contributions of the research nurses in Shetland, the administrative team in Edinburgh, the people of Shetland, and the data analysts in particular Dr Thibaud Boutin for the genotype imputation to the HRC reference panel and Dr Peter Joshi for setting up the GWAS analysis pipeline used in this analysis.

## UConn Health research collaboration

We like to thank Dr. J. L. Cotney (Genetics & Genome Sciences, UConn Health) for investigating the association of craniofacial eQTL and mQTL with our HC GWAS hits.

## Research Centre Jülich research collaboration

We would like to thank Dr. T.W. Mühleisen and Prof. S. Cichon (Department of Biomedicine, University Hospital Basel) for discussion of the functionalities of our HC GWAS hits.

## The Cohorts for Heart and Aging Research in Genomic Epidemiology (CHARGE) Consortium

We would like to thank the CHARGE consortium for making their summary statistics publicly available, and therefore for us to incorporate this information into our analysis.

## The Enhancing NeuroImaging Genetics through Meta-Analysis (ENIGMA) Consortium

We have incorporated the summary statistics from ENIGMA Consortium into analysis of this paper, therefore we gratefully acknowledge their efforts and making their data publicly available.

## Author contributions

B. S.P, V.V, E.Z, G.D, V.J, C.E.P, K.B, H.B, and D.M designed and supervised the research. S.H, C.Y.S, C. H, B.P, J.F, M.C.M-G, C.W, T.S.A, M B, D.F, L.S, I T, K.W, A.J, L.C, J.H, and J.L.M analyzed genetic data. V.I, F.R and T.J.C provided methodological support. G.D.S and S.E.F contributed ideas in the initial stage of the project. B.S.P, S.H and C.Y.S wrote the manuscript. All authors read and commented on the manuscript.

## References

1. Fabbri, M., et al, The skull roof tracks the brain during the evolution and development of reptiles including birds. Nature ecology & evolution, 2017. 1(10): p. 1543.

2. Koyabu, D., et al, Mammalian skull heterochrony reveals modular evolution and a link between cranial development and brain size. Nature communications, 2014. 5: p. 3625.

3. Harris, S.R., Measuring head circumference: Update on infant microcephaly. Canadian Family Physician, 2015. 61(8): p. 680–684.

4. Maunu, J., et al, Brain and ventricles in very low birth weight infants at term: a comparison among head circumference, ultrasound, and magnetic resonance imaging. Pediatrics, 2009. 123(2): p. 617–626.

5. Bartholomeusz, H., E. Courchesne, and C. Karns, Relationship between head circumference and brain volume in healthy normal toddlers, children, and adults. Neuropediatrics, 2002. 33(05): p. 239–241.

6. De Onis, M., et al, Les standards de croissance de l’Organisation mondiale de la sante pour les nourrissons et les jeunes enfants. Archives de pédiatrie, 2009. 16(1): p. 47–53.

7. Cole, T.J., J.V. Freeman, and M.A. Preece, British 1990 growth reference centiles for weight, height, body mass index and head circumference fitted by maximum penalized likelihood. Statistics in medicine, 1998. 17(4): p. 407–429.

8. Scheffler, C., H. Greil, and M. Hermanussen, The association between weight, height, and head circumference reconsidered. Pediatric research, 2017. 81(5): p. 825–830.

9. Hshieh, T.T., et al, Head circumference as a useful surrogate for intracranial volume in older adults. International psychogeriatrics, 2016. 28(1): p. 157–162.

10. Smit, D.J., et al, Heritability of head size in Dutch and Australian twin families at ages 0-50 years. Twin Research and Human Genetics, 2010. 13(4): p. 370–380.

11. Richtsmeier, J.T. and K. Flaherty, Hand in glove: brain and skull in development and dysmorphogenesis. Acta neuropathologica, 2013. 125(4): p. 469–489.

12. Adams, H.H., et al, Novel genetic loci underlying human intracranial volume identified through genome-wide association. Nature neuroscience, 2016. 19: p. 1569–1582.

13. Taal, H.R., et al, Common variants at 12q15 and 12q24 are associated with infant head circumference. Nature genetics, 2012. 44(5): p. 532–538.

14. Huang, J., et al, Improved imputation of low-frequency and rare variants using the UK10K haplotype reference panel. Nature communications, 2015. 6.

15. Consortium, H.R., A reference panel of 64,976 haplotypes for genotype imputation. Nature genetics, 2016. 48(10): p. 1279–1283.

16. Manousaki, D., et al, Low-frequency synonymous coding variation in CYP2R1 has large effects on vitamin D levels and risk of multiple sclerosis. The American Journal of Human Genetics, 2017. 101(2): p. 227–238.

17. Tachmazidou, I., et al, Whole-genome sequencing coupled to imputation discovers genetic signals for anthropometric traits. The American Journal of Human Genetics, 2017. 100(6): p. 865–884.

18. Cole, T.J., M.D. Donaldson, and Y. Ben-Shlomo, SITAR—a useful instrument for growth curve analysis. International journal of epidemiology, 2010. 39(6): p. 1558–1566.

19. Wood, A.R., et al, Defining the role of common variation in the genomic and biological architecture of adult human height. Nature genetics, 2014. 46(11): p. 1173–1186.

20. de Leeuw, C.A., et al, MAGMA: generalized gene-set analysis of GWAS data. PLoS computational biology, 2015. 11(4): p. e1004219.

21. Barbeira, A.N., et al, Exploring the phenotypic consequences of tissue specific gene expression variation inferred from GWAS summary statistics. bioRxiv, 2017.

22. Bulik-Sullivan, B.K., et al, LD Score regression distinguishes confounding from polygenicity in genome-wide association studies. Nature genetics, 2015. 47(3): p. 291–295.

23. Bulik-Sullivan, B., et al, An atlas of genetic correlations across human diseases and traits. Nature genetics, 2015. 47(11): p. 1236–1241.

24. St Pourcain, B., et al, Developmental Changes Within the Genetic Architecture of Social Communication Behavior: A Multivariate Study of Genetic Variance in Unrelated Individuals. Biological psychiatry, 2017.

25. Davydov, E.V., et al, Identifying a high fraction of the human genome to be under selective constraint using GERP++. PLoS computational biology, 2010. 6(12): p. e1001025.

26. Watanabe, K., et al, Functional mapping and annotation of genetic associations with FUMA. Nature communications, 2017. 8(1): p. 1826.

27. Ng, B., et al, An xQTL map integrates the genetic architecture of the human brain’s transcriptome and epigenome. Nature neuroscience, 2017. 20(10): p. 1418.

28. Wilderman, A., et al, High Resolution Epigenomic Atlas of Early Human Craniofacial Development. bioRxiv, 2017: p. 135368.

29. Millard, L.A., et al, Software Application Profile: PHESANT: a tool for performing automated phenome scans in UK Biobank. International Journal of Epidemiology, 2017.

30. Vousden, K.H. and C. Prives, Blinded by the light: the growing complexity of p53. Cell, 2009. 137(3): p. 413–431.

31. Olivier, M. and P. Hainaut, IARC TP53 Database, in Encyclopedia of Cancer, M. Schwab, Editor. 2011, Springer Berlin Heidelberg: Berlin, Heidelberg. p. 1799–1802.

32. Stacey, S.N., et al, A germline variant in the TP53 polyadenylation signal confers cancer susceptibility. Nature genetics, 2011. 43(11): p. 1098–1103.

33. Khoury, M.P. and J.-C. Bourdon, The isoforms of the p53 protein. Cold Spring Harbor perspectives in biology, 2010. 2(3): p. a000927.

34. Khoury, M.P. and J.-C. Bourdon, p53 isoforms: an intracellular microprocessor? Genes & cancer, 2011. 2(4): p. 453–465.

35. Melin, B.S., et al, Genome-wide association study of glioma subtypes identifies specific differences in genetic susceptibility to glioblastoma and non-glioblastoma tumors. Nature genetics, 2017. 49(5): p. 789.

36. Diskin, S.J., et al, Rare variants in TP53 and susceptibility to neuroblastoma. Journal of the National Cancer Institute, 2014. 106(4): p. dju047.

37. Trampush, J., et al, GWAS meta-analysis reveals novel loci and genetic correlates for general cognitive function: a report from the COGENT consortium. Molecular psychiatry, 2017. 22(3): p. 336–345.

38. Rinon, A., et al, p53 coordinates cranial neural crest cell growth and epithelial-mesenchymal transition/delamination processes. Development, 2011. 138(9): p. 1827–1838.

39. Jones, N.C., et al, Prevention of the neurocristopathy Treacher Collins syndrome through inhibition of p53 function. Nature medicine, 2008. 14(2): p. 125–133.

40. Jin, S.-W., K.-B. Sim, and S.-D. Kim, Development and growth of the normal cranial vault: an embryologic review. Journal of Korean Neurosurgical Society, 2016. 59(3): p. 192.

41. Kim, N.H., et al, p53 and microRNA-34 are suppressors of canonical Wnt signaling. Sci. Signal., 2011. 4(197): p. ra71–ra71.

42. Mulligan, K.A. and B.N. Cheyette, Wnt signaling in vertebrate neural development and function. Journal of Neuroimmune Pharmacology, 2012. 7(4): p. 774–787.

43. Heydeck, W. and A. Liu, PCP effector proteins inturned and fuzzy play nonredundant roles in the patterning but not convergent extension of mammalian neural tube. Developmental Dynamics, 2011. 240(8): p. 1938–1948.

44. Mirzaa, G.M., et al, De novo CCND2 mutations leading to stabilization of cyclin D2 cause megalencephaly-polymicrogyria-polydactyly-hydrocephalus syndrome. Nature genetics, 2014. 46(5): p. 510.

45. Rivière, J.-B., et al, De novo germline and postzygotic mutations in AKT3, PIK3R2 and PIK3CA cause a spectrum of related megalencephaly syndromes. Nature genetics, 2012. 44(8): p. 934.

46. Alcantara, D., et al, Mutations of AKT3 are associated with a wide spectrum of developmental disorders including extreme megalencephaly. Brain, 2017. 140(10): p. 2610–2622.

47. Neubauer, S. and J.-J. Hublin, The evolution of human brain development. Evolutionary Biology, 2012. 39(4): p. 568–586.

48. Hagenaars, S.P., et al, Shared genetic aetiology between cognitive functions and physical and mental health in UK Biobank (N= 112 151) and 24 GWAS consortia. Molecular psychiatry, 2016. 21(11): p. 1624.

49. Martini, M., et al, Head circumference-a useful single parameter for skull volume development in cranial growth analysis? Head & face medicine, 2018. 14(1): p. 3.

50. Turley, P., et al, Multi-trait analysis of genome-wide association summary statistics using MTAG. Nature genetics, 2018: p. 1.

51. Zheng, H.-F., et al, Whole-genome sequencing identifies EN1 as a determinant of bone density and fracture. Nature, 2015. 526(7571): p. 112.

52. Consortium, U.K., The UK10K project identifies rare variants in health and disease. Nature, 2015. 526(7571): p. 82–90.

53. Winkler, T.W., et al, Quality control and conduct of genome-wide association meta-analyses. Nature protocols, 2014. 9(5): p. 1192.

54. Yang, J., et al, GCTA: a tool for genome-wide complex trait analysis. The American Journal of Human Genetics, 2011. 88(1): p. 76–82.

55. Li, M.-X., et al, Evaluating the effective numbers of independent tests and significant p-value thresholds in commercial genotyping arrays and public imputation reference datasets. Human genetics, 2012. 131(5): p. 747–756.

56. Marchini, J., et al, A new multipoint method for genome-wide association studies by imputation of genotypes. Nature genetics, 2007. 39(7): p. 906.

57. Zhou, X. and M. Stephens, Genome-wide efficient mixed-model analysis for association studies. Nature genetics, 2012. 44(7): p. 821.

58. Willer, C.J., Y. Li, and G.R. Abecasis, METAL: fast and efficient meta-analysis of genomewide association scans. Bioinformatics, 2010. 26(17): p. 2190–2191.

59. Stein, J.L., et al, Identification of common variants associated with human hippocampal and intracranial volumes. Nature genetics, 2012. 44(5): p. 552–561.

60. Consortium, E.G.G., Common variants at 6q22 and 17q21 are associated with intracranial volume. Nature genetics, 2012. 44(5): p. 539–544.

61. Hibar, D.P., et al, Common genetic variants influence human subcortical brain structures. Nature, 2015. 520(7546): p. 224–229.

62. Elliott, L., et al, The genetic basis of human brain structure and function: 1,262 genome-wide associations found from 3,144 GWAS of multimodal brain imaging phenotypes from 9,707 UK Biobank participants. bioRxiv, 2017.

63. Rietveld, C.A., et al, GWAS of 126,559 individuals identifies genetic variants associated with educational attainment. science, 2013. 340(6139): p. 1467–1471.

64. Zheng, J., et al, LD Hub: a centralized database and web interface to perform LD score regression that maximizes the potential of summary level GWAS data for SNP heritability and genetic correlation analysis. Bioinformatics, 2017. 33(2): p. 272–279.

65. Finucane, H., et al, Heritability enrichment of specifically expressed genes identifies diseaserelevant tissues and cell types. bioRxiv, 2017: p. 103069.

66. Sudlow, C., et al, UK biobank: an open access resource for identifying the causes of a wide range of complex diseases of middle and old age. PLoS medicine, 2015. 12(3): p. e1001779.

